# Small RNAs regulation and genomic harmony: insights into allopolyploid evolution in marsh orchids (*Dactylorhiza*)

**DOI:** 10.1101/2024.11.29.626004

**Authors:** Mimmi C. Eriksson, Matthew Thornton, Emiliano Trucchi, Thomas M. Wolfe, Francisco Balao, Mikael Hedrén, Ovidiu Paun

## Abstract

Hybridization and polyploidy are major drivers of plant diversification, often accompanied by shifts in gene expression and genome composition. Small RNAs (smRNAs) are thought to influence such genomic changes, particularly through their interactions with transposable elements (TEs).
We comparatively quantified smRNAs in established sibling allopolyploids *Dactylorhiza majalis* and *D. traunsteineri* and their diploid progenitors to assess how independent allopolyploidization events shaped smRNA landscapes.
Despite independent origins, the allotetraploids exhibited substantial overlap in smRNA composition, including transgressive accumulation of smRNA near genes related to transcriptional regulation, cell division and stress response. Consistently, TE-associated 24 nt smRNAs more closely resembled the paternal and larger genome, while shorter smRNAs typically reflected the maternal and smaller genome. Nevertheless, distinct patterns were also evident: the older *D. majalis* showed greater accumulation of smRNAs near genes involved in transcriptional and translational regulation, while the younger *D. traunsteineri* displayed stronger non-additive patterns, suggesting ongoing resolution of post-polyploid meiotic and mitotic instability.
Our results reveal both convergence and divergence in smRNA landscapes among independently formed allopolyploids. Our study highlights the central role of smRNAs in resolving genomic conflict, with possible implications for functional divergence and ecological innovation during polyploid evolution.

**Plain-language summary:** We studied two sibling marsh orchid species that arose through hybridization and genome doubling. By comparing their small RNA molecules and how these associate with genes, we found both shared and species-specific patterns. These differences reflect how the orchids’ genomes have changed since formation, and may explain how repeated polyploidy events contribute to genetic and ecological diversity in plants.

## INTRODUCTION

Allopolyploidization—the combination of whole-genome duplication (WGD) and interspecific hybridization—can be a potent evolutionary force, generating genetic diversity, gene redundancy, and buffering capacity that may contribute to evolutionary innovation and ecological diversification. However, the initial merger of divergent genomes into a single nucleus can destabilize genome regulation, triggering “genomic shock” (McClintock, 1984), on the background of a severe genetic bottleneck.

Small noncoding RNAs (smRNAs) have emerged as key mediators during post-polyploidization genome stabilization (e.g., Ha *et al*., 2009; Shen *et al*., 2017; Cavé-Radet *et al*., 2020). These short RNA molecules suppress transposable elements (TEs) and fine-tune gene expression, promoting both genome integrity and adaptive responses. Specifically, repeat-associated or heterochromatic small-interfering RNAs (ra-siRNAs/het-siRNAs; Shen *et al*., 2017) help maintain chromatin stability, while microRNAs (miRNAs) can modulate gene expression with phenotypic implications (Ha *et al*., 2009; Ghani *et al*., 2014; Jiang *et al*., 2021). In addition, plant smRNAs orchestrate abiotic and biotic stress responses (Kaur *et al*., 2021; Tiwari & Rajam, 2022; Zhan & Meyers, 2024), potentially facilitating adaptation to novel, non-parental environments. These dual roles make smRNAs integral to allopolyploid success.

In general, smRNAs operate through three main mechanisms: (1) post-transcriptional gene silencing (PTGS), binding to RNAs to repress translation or promote cleavage and degradation; (2) transcriptional gene silencing (TGS), directing epigenetic silencing at homologous genomic loci; and (3) splicing regulation, interacting with pre-mRNAs to influence splice-site selection (Borges & Martienssen, 2015; Wang & Chekanova, 2016). In plants, smRNAs are classified by biogenesis into miRNAs (20–22 nt long, hairpin-derived), and small-interfering RNAs (siRNAs, from double stranded RNA) (Borges & Martienssen, 2015; Chen & Rechavi, 2021), with siRNAs further divided into: (*i*) ra-siRNAs (24 nt and primarily involved in TGS), (*ii*) secondary siRNAs (∼21–22nt long, which can function in both TGS and PTGS pathways), and (*iii*) natural antisense transcript siRNA (nat-siRNA, 21–24 nt, acting both in *cis* and *trans* via PTGS) (Borges & Martienssen, 2015; Chen a& Rechavi, 2021).

Recent reviews (e.g., Ariel & Manavella, 2021; Tossolini *et al*., 2025) highlight the pivotal link between TE mobilization and smRNA evolution. TEs not only contribute regulatory capacity through enhancer-derived transcripts (e.g., in wheat; TE-initiated enhancer-like elements, Xie *et al*., 2023), but also spur the emergence of smRNAs and long non-coding RNAs that diversify gene regulation and support genome plasticity (Hassan *et al*., 2024; Tossolini *et al*., 2025).

WGD and environmental stress are known to lead to TE activation, resulting in accumulation of repeat-derived smRNAs, boosting both TGS and PTGS pathways (Ito *et al*., 2011; Kenan-Eichler *et al*., 2011; Marí-Ordóñez *et al*., 2013; Fultz *et al*., 2015; Xiao & Wang, 2025). For example, resynthesized *Arabidopsis* allotetraploids rapidly restore ra-siRNA populations after WGD, while miRNAs show non-additive expression patterns and play key roles in stabilizing hybrid genomes (Ha *et al*., 2009). Similarly, early generations *Brassica napus* allotetraploids exhibit a surge in TE-derived 21 nt siRNAs, especially from Gypsy elements, helping to silence newly active TEs (Martinez Palacios *et al*., 2019). Cavé-Radet *et al*. (2020, 2023) observed stronger miRNA responses in allopolyploid *Spartina* compared to homoploid hybrids, suggesting a more pronounced regulatory impact of WGD than hybridization alone (Cavé-Radet *et al*., 2020). On the other hand, the divergence between progenitors determines the extent of genomic shock in wheat allotetraploids, including enhanced smRNA responses associated with TEs in hybrid crosses between parents with differing TE load, compared to those with similar TE content (Jiao *et al*., 2018).

Despite growing interest in the role of smRNAs in allopolyploid evolution, most research has focused on synthetic or early-generation polyploids. To our knowledge, only Cavé-Radet *et al*., (2020) have examined smRNAs in wild allopolyploids, albeit also in early stages (*Spartina* F_₁_ hybrids and *S. anglica*, ∼170 years old). Thus, the long-term dynamics, regulatory roles and potential ecological relevance of smRNAs in more established allopolyploid lineages remain poorly explored.

In this study, we focus on two ecologically-distinct, established, sibling orchid allotetraploids, *Dactylorhiza majalis s.str.* (Rchb.) P.F.Hunt & Summerh. and *D. traunsteineri* (Saut. ex Rchb.) Soó, and their parents, to address the long-term fate and potential roles of smRNAs following allopolyploidization. The marsh orchids within *D. majalis s.l.* include several independently formed sibling allotetraploids, each likely with multiple origins involving overlapping stocks of parental populations (Brandrud *et al*., 2020). The focal allotetraploids here share a complex history of repeated polyploidization and/or hybridization and back-crossing, but are today reasonably well separated in morphology and habitat preferences (Fig. 1a; Wolfe *et al*., 2023). They are estimated to have formed approximately 104k (*D. majalis*) and 74k (*D. traunsteineri*) generations ago (Hawranek, 2021), from the diploid progenitors *D. fuchsii* (Druce) Soó (maternal, smaller genome) and *D. incarnata* (L.) Soó (paternal, larger genome) (Fig. 1a) (Aagard *et al*., 2005; Brandrud *et al*., 2020; Eriksson *et al*., 2022). Transcriptomic comparisons of the diploid progenitors have revealed large-scale gene expression differences, importantly, including differences in smRNAs associated with TEs and PTGS pathways (Balao *et al*., 2017). Further, Eriksson *et al*. (2022) reported significant differences in both genome size and TE load between the two diploid progenitors.

**Figure 1.**
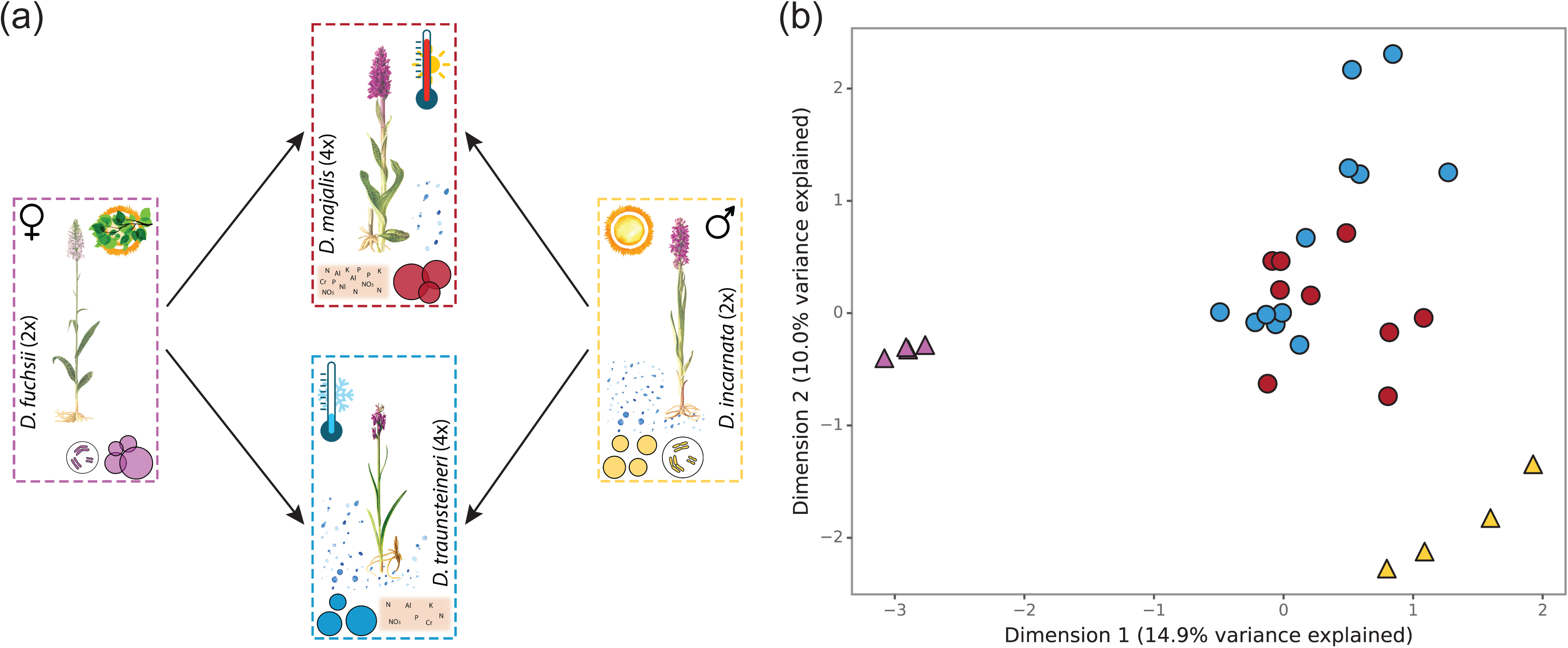
Study system comprising two diploids (*Dactylorhiza fuchsii* and *D. incarnata*) and their sibling allotetraploids (*D majalis* and *D. traunsteineri*). **(a)** Overview of the four species. Coloured circles and their relative positions illustrate population densities. Cell icons indicate difference in genome size between the diploids, while gender symbols show the direction of hybridization leading to the formation of the allopolyploids. Additional symbols denote broad ecological preferences. **(b)** Multidimensional scaling (MDS) plot based on small RNA count data. Colours correspond to the species frames shown in (a); triangles indicate diploid accessions, and circles, allotetraploid accessions.

Distinct ecological, spatial, morphological and physiological properties have been reported for the two sibling allopolyploids (Paun *et al*., 2010, 2011; Balao *et al*., 2016; Wolfe *et al*., 2023)*. D. majalis* is distributed across continental Europe, ranging from the Pyrenees to Scandinavia, with a broader ecological niche (i.e., mesic to moist meadows) and less geographically-driven genetic differentiation than *D. traunsteineri* which is found in disjunct marshes and fens in the Alps, Scandinavia and British Isles. Wolfe and colleagues (2023) reported physiological differences in photosynthesis and nutrient transport consistent with the contrasting ecological conditions preferred by the two allotetraploids. These appear to have helped the two allotetraploids adapt to different soil chemistry conditions, with *D. traunsteineri* restricted to wet, nutrient-poor soils, in particular with extremely low levels of available nitrate (Wolfe *et al*., 2023). Furthermore, reciprocal transplant experiments have shown that much of the gene expression divergence between the sibling allopolyploids is environmentally plastic, and appeared to be associated to distinct, environmentally-driven, biotic communities in their roots (Emelianova *et al*., 2025). In stark contrast, analysis of the genomes of several *Dactylorhiza* allopolyploids found relatively little difference between *D. majalis* and *D. traunsteineri* both in genome size and TE activity, suggesting a strong control of TEs following recurrent allopolyploidizations (Eriksson *et al*., 2022).

In this study, we specifically investigate: (1) how smRNA abundance in the allopolyploids reflects parental patterns—whether it follows additive, dominant, or transgressive trends— and how consistent these patterns are across the sibling allopolyploids; (2) the extent and similarity of smRNA association with TEs in the two sibling allotetraploids; and (3) how variation in smRNA patterns may relate to distinct eco-physiological traits of *D. majalis* and *D. traunsteineri*.

## MATERIALS AND METHODS

### Sampling

Adult specimens of each species of interest were transplanted from multiple localities across Europe to a common garden in Vienna, Austria. *Dactylorhiza* includes perennial plants that produce a new tuber yearly, which supports growth in the following year. Two years after transplantation, in the morning of 14 May 2014, leaf tissue at a similar developmental stage for all accessions was fixed in RNAlater (Sigma) and stored at -80°C until further processing. Our final sampling included four accessions of each diploid (*D. incarnata* and *D. fuchsii*), together with nine (*D. majalis*) and twelve (*D. traunsteineri*) individuals of each allotetraploid (Supplementary Table S1). With few exceptions, each accession of each species originated from different localities across Europe. The diploid smRNA data has been previously investigated by Balao *et al*. (2017). Independent RNA isolations from the same tissue fixations of *D. majalis* and *D. traunsteineri* were used for RNA-seq analyses by Wolfe *et al*. (2023).

### Library preparations and sequencing

Small RNA extraction, library preparation and sequencing for this experiment was previously described for diploid investigations (Balao *et al*., 2017). In short, isolation of smRNA was performed with the mirVana miRNA Isolation Kit (Life Technologies). The smRNA-enriched RNA extracts were further purified by gel size selection on 15% TBE-Urea pre-casted gels (Life Technologies). The small RNA libraries were prepared with the NEBNext Multiplex Small RNA Library Prep Set for Illumina (New England Biolabs). Illumina sequencing as 50 bp single-end reads was performed at the Vienna BioCenter Core Facilities (VBCF; https://www.viennabiocenter.org/).

### Read processing and quantifying

The libraries were demultiplexed using BamIndexDecoder v.1.03 (http://wtsi-npg.github.io/illumina2bam/#BamIndexDecoder). Raw reads were adapter-trimmed, size-selected for the 20-24 nt range, and then quality filtered using CLC Genomics Workbench v.8.0 (QIAGEN). Filtered reads were mapped to the *Dactylorhiza incarnata* reference genome v.1.0 (Wolfe *et al*., 2023) with STAR v.2.7.3a (Dobin *et al*., 2013), using -- alignIntronMax 1 and --outFilterMismatchNoverLmax 0.05. A single-pass alignment approach was used, retaining multi-mapping reads below a threshold of 100 (-- outFilterMulitmapNmax 100).

An annotation-free approach was initially used to identify genomic regions (genic and non-genic) that were potentially “targeted” by smRNAs, defined here as regions showing sequence homology with smRNAs and therefore indicating an association between smRNAs and the corresponding genomic sequence. Throughout this study, we use the term *“targeting”* in a purely operational sense to describe genomic regions showing sequence complementarity to smRNAs. This definition does not assume a direct causal or regulatory interaction between smRNAs and the corresponding genomic loci. Rather, it denotes potential sites of smRNA association inferred from sequence homology. While such associations often coincide with transcriptionally or post-transcriptionally regulated regions (e.g. in the case of heterochromatic siRNAs or miRNAs; Borges & Martienssen, 2015), they may also reflect indirect or distal effects, or even passive sequence associations without immediate regulatory function.

After sorting bam files by reference coordinates with Samtools v.1.10 (Danacek *et al*., 2011), smRNA read coverage was assessed for each individual in non-overlapping 100 bp windows across the genome using bamCoverage from deepTools2 v.3.5.1 (Ramirez *et al*., 2016). Windows with a minimum coverage of ten reads in at least one sample were retained in a bed file, and adjacent windows were merged using bedtools merge v.2.29.2 (Quinlan & Hall, 2010). The resulting bed file of retained regions (i.e., peaks of smRNA “targeting”) was converted to gff3 format using a custom python script. Counts were then obtained with featureCounts from Subread v.2.0.0 (Liao *et al*., 2014), accounting for multiple mappers fractionally (-M --fraction), to include potential trans-associations. The full pipeline, scripts, raw count file, and peak gff3 file are available at https://github.com/mc-er/dact-sRNAs.

### Differential smRNA “targeting” analysis

Differently “targeted” (DT) regions were identified following a similar approach to that used for detecting differentially expressed genes, using counts of smRNAs at peaks as input. Analyses were performed in R v.4.1.1 and the Bioconductor package EdgeR v.3.34.0 (Robinson *et al*., 2010; McCarthy *et al*., 2012). The count matrix obtained from featureCounts was filtered to retain genomic regions with a minimum smRNA mapping of one count per million (CPM) in at least 25% of samples within a group and a minimum of two samples overall. To account for differences in library size, data were normalised using the weighted trimmed mean of M-values (TMM) method. A quasi-likelihood negative binomial model was applied to identify DT regions across the genome, comparing either (i) the two sibling allotetraploids or (ii) each allotetraploid to its parental diploid. P-values were corrected for multiple testing using the Benjamini–Hochberg (BH) procedure, and only genomic regions passing an FDR < 0.05 were kept for further analysis.

Analyses of genomic regions around TEs were performed separately for reads of 20–23 nt, and for 24 nt reads (the latter representing potentially heterochromatic smRNAs, hetsiRNAs; Borges & Martienssen, 2015). For analyses of genic regions, all read lengths (20–24 nt) were combined into a single analysis per comparison.

### Assessing genomic interactions

Differentially targeted peaks passing an FDR threshold of 0.05 were classified into three categories of genomic interactions (Supplementary Table S2): (1) additivity—the polyploid showed lower targeting relative to one diploid and higher targeting relative to the other, (2) transgressive—the polyploid showed increased or decreased targeting relative to both diploids, and (3) dominance—the polyploid showed no significant difference relative to one diploid (the “dominant” parent) but differed significantly relative to the other diploid (illustrated on the Y-axis of Fig. 4c-f). Deviation from the stochastically expected number of shared peaks between the two tetraploids in each category was tested using a hypergeometric test implemented in the Bioconductor package SuperExactTest (Wang *et al*., 2015).

### Functional interpretation of differentially targeted genes

To gain insights into potential roles for smRNA associations in shaping the divergent ecological properties of the sibling allopolyploids, we performed gene ontology (GO) enrichment analysis of DT genes. Because the molecular consequences of smRNA association depend on the location of the targeted region, we analysed regions around promoters separately from those within exons or introns. Genes associated with smRNAs around promoters would be expected to be transcriptionally down-regulated or silenced, whereas associations within gene bodies may indicate post-transcriptional effects or altered splicing. GO enrichment analyses for genes potentially affected by DT peaks were performed in R v.4.1.1 using the Bioconductor package topGO v.2.44.0 (Alexa & Rahnenfuhre, 2021). Fisher’s exact tests were run using the “weight01” algorithm to test for over-representation of specific functional categories. P-values were adjusted for multiple testing using the FDR method, with significance set at FDR < 0.05.

## RESULTS

### smRNAs and their genomic targets

Sequencing produced an average 13.5 M reads per accession (±5.9 M SD). After size selection and quality filtering, an average of 5.8 M reads (±3.2 M SD) of 20–24 nt were retained (Supplementary Table S3). On average, 83.1% (±4.2% SD) of these reads mapped to the *D. incarnata* reference genome, with no apparent mapping bias across species (Supplementary Table S3). Read length distributions matched the expected profile for smRNAs (Borges & Martienssen, 2015), with modes at 21–22 nt and 24 nt in all species (Fig. 2a).

**Figure 2.**
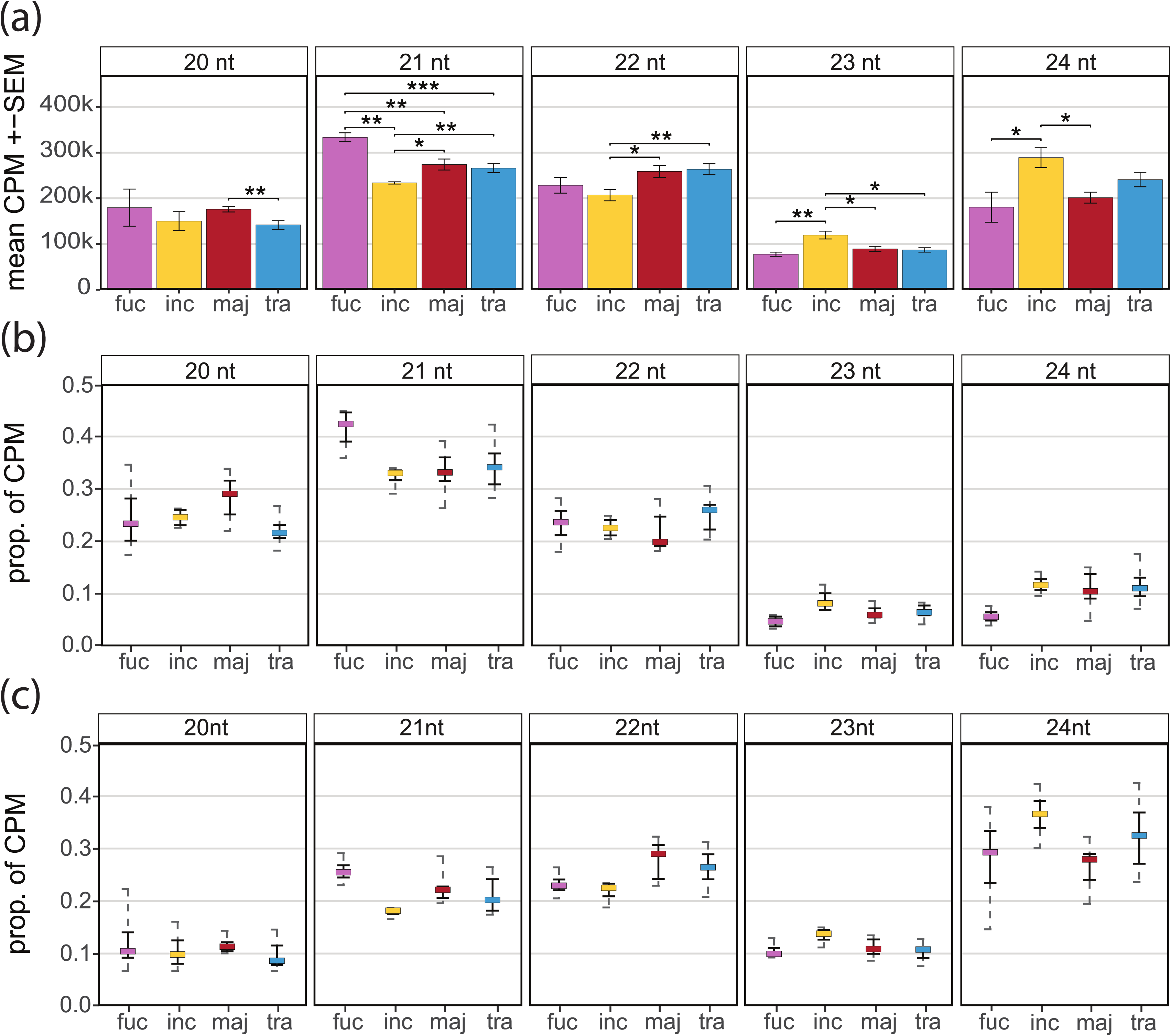
Small RNA read length distributions across species. **(a)** Mean counts per million (CPM) for each read length; error bars represent the standard error of the mean. Horizontal lines indicate significant differences between species, with asterisks denoting significance levels: * P < 0.05, ** P < 0.01, *** P < 0.001. **(b, c)** Proportions of reads by species and read length mapped to genomic features: (b) exons and (c) transposable elements. fuc*, D. fuchsii*; inc, *D. incarnata*; maj, *D. majalis*; tra, *D. traunsteineri*.

*D. fuchsii* exhibited higher CPM of 20–22 nt smRNAs compared to *D. incarnata*, with a significant difference observed only for 21 nt smRNAs (Fig 2a–c). The reverse trend was found for the 23–24 nt smRNAs, which were more abundant in *D. incarnata* (Fig. 2a). The two allotetraploids were more similar to each other than the diploids (Fig. 2a). For 21 nt smRNAs, both allotetraploids displayed significantly higher CPM than the paternal species (*D. incarnata*), but significantly lower CPM than the maternal species (*D. fuchsii*) (Fig. 2a). A large proportion of 20–22 nt smRNAs mapped to exonic regions, while 24 nt smRNAs predominantly mapped to TEs and intergenic regions (Fig. 2b–c, Supplementary Fig. S1).

Our annotation-free approach identified 2,454,439 putative smRNA peaks across the reference genome. After filtering for CPM > 1 in at least two samples and 25% of samples per group, between 102,222 and 370,440 peaks were retained depending on the comparison. A multidimensional scaling (MDS) plot based on all 20–24 nt smRNAs across 212,434 peaks clearly separated the two diploid species, and distinguished diploids from allotetraploids, with the latter overlapping centrally (Fig. 1b). Within *D. traunsteineri* a geographical clustering was apparent (not shown).

### smRNA patterns in ecologically divergent allopolyploids

We compared the two sibling allotetraploids using the combined 20–24 nt reads, retaining 152,503 peaks passing the CPM threshold (Supplementary Table S4). Among these, 2,106 DT peaks were identified between *D. traunsteineri* and *D. majalis*, of which approximately half (1,016 peaks) were associated with genes. Gene-associated peaks were defined as those overlapping a gene model (exons or introns) or within 1,000 bp upstream of a gene model (hereafter referred to as ‘promoters’; Supplementary Table S4).

Most genic DT peaks (762) were located within introns, and 82 overlapped multiple genic features (intron, exon, and/or promoters) (Supplementary Table S4). Across all genic categories, the older allotetraploid *D. majalis* displayed approximately three times as many peaks with high smRNA targeting compared to *D. traunsteineri* (Supplementary Table S4, Fig. 3a–c).

**Figure 3.**
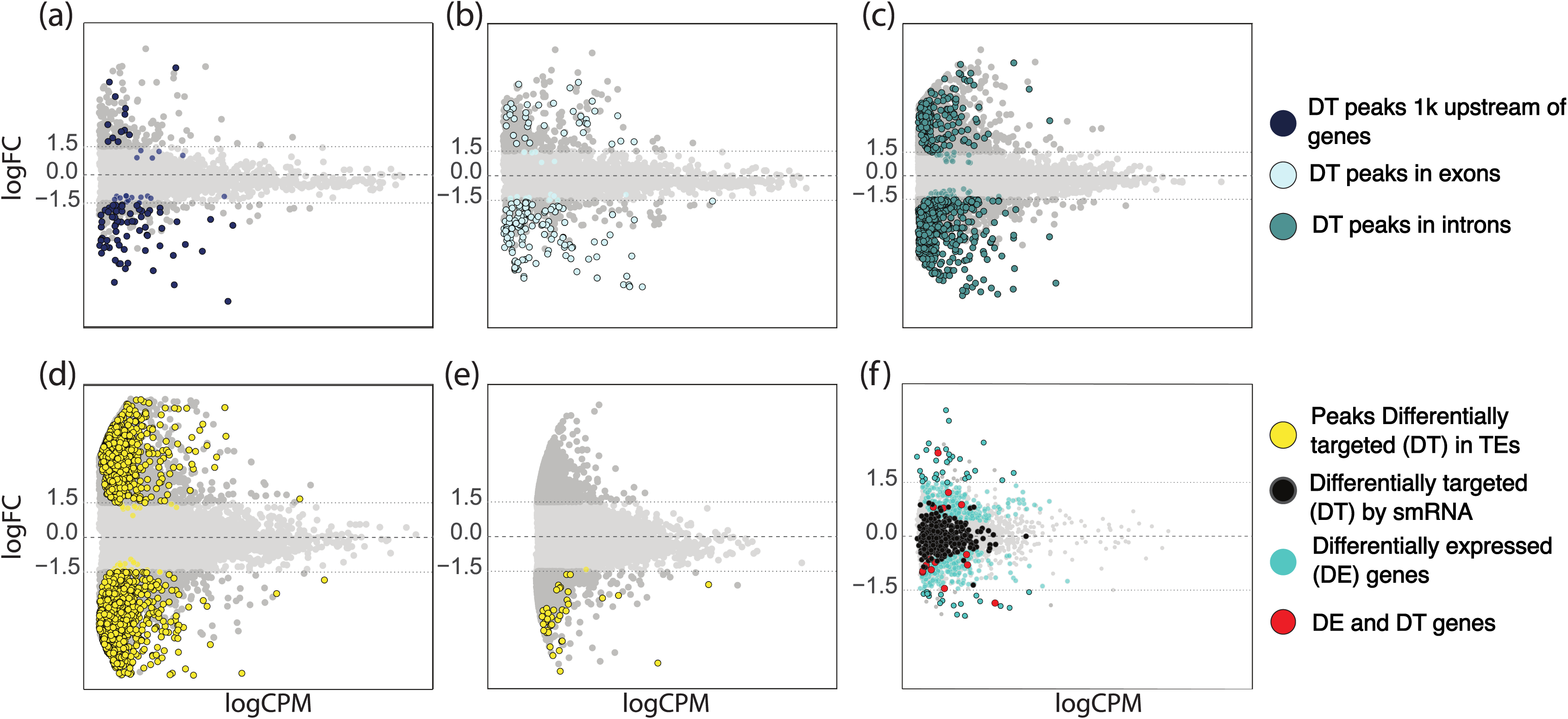
MA plots showing differences in small RNA (smRNA) “targeting” between *Dactylorhiza majalis* and *D. traunsteineri*. The central dashed line marks logFC = 0, while dotted lines indicate logFC = -1.5 and 1.5. Negative logFC values represent higher smRNA targeting in *D. majalis* relative to *D. traunsteineri*, whereas positive logFC values indicate higher targeting in *D. traunsteineri*. **(a–c)** Peaks differently targeted in genic regions by 20– 24 nt smRNAs: **(a)** promoter/1,000 bp upstream gene models, **(b)** exons and **(c)** introns. **(d)** Transposable elements (TEs) targeted by 20–23 nt smRNAs. **(e)** TEs targeted by 24 nt smRNAs. **(f)** MA plot of RNA-seq data from Wolfe *et al*. (2023), showing genes differentially expressed (DE) and/or DT by smRNAs as indicated in the legend.

Comparison of DT peaks with previously identified differentially expressed (DE) genes from the same experiment (Wolfe *et al*., 2023) revealed that most genes with DT peaks did not show differential expression between the two allopolyploids, consistent with potential post-transcritional effects or alternative splicing rather than direct transcriptional regulation (Fig. 3f, Supplementary Table S5).

GO enrichment analyses of genes associated with DT peaks between the allotetraploids revealed more enriched terms in *D. majalis* than *D. traunsteineri* (Fig. 5a, Supplementary Table S6). Most enriched terms were for genes targeted within gene bodies and related to transcription/translation regulation, transmembrane transport, response to biotic and abiotic stress, DNA repair, and cell cycle control (Supplementary Table S6).

### smRNA targeting and genomic interactions

We further compared 20-24 nt smRNA patterns of each allotetraploid to those of the parental diploids. After filtering, 212,434 peaks were retained (Supplementary Table S4). Both allotetraploids showed broadly similar targeting patterns relative to the diploids, although the younger *D. traunsteineri* displayed greater divergence than *D. majalis* (Supplementary Table S4, Supplementary Fig. S2). Allotetraploids showed larger differences from the maternal parent (*D. fuchsii*) than from the paternal *D. incarnata* (Supplementary Table S4, Supplementary Fig. S2).

DT peaks were classified into four interaction types: additive, transgressive, and dominance towards either diploid. The majority of peaks showed no change relative to either diploid. Among DT peaks, non-additive patterns predominated, with dominance most frequent, particularly towards *D. incarnata* (Supplementary Table S7, Fig. 4a–f, Supplementary Fig. S3–S4). Notably, in intergenic regions lacking annotated TEs, a higher proportion of smRNA peaks showing dominance towards *D. fuchsii* than in other genomic contexts (Supplementary Table S7, Fig. 4d, Supplementary Fig. S3). Among transgressive peaks, both allotetraploids most often displayed higher smRNA targeting relative to both diploids (Supplementary Table S7, Fig. 4a, Fig. 4c–f, Supplementary Fig. S3–S4). A notable general pattern across interaction types was that a large proportion of DT peaks were shared between the sibling allotetraploids (Supplementary Table S7, Fig. 4b–c, Supplementary Fig. S3–S4), despite their independent origins (Brandrud *et al*., 2020; Hawranek, 2021).

**Figure 4.**
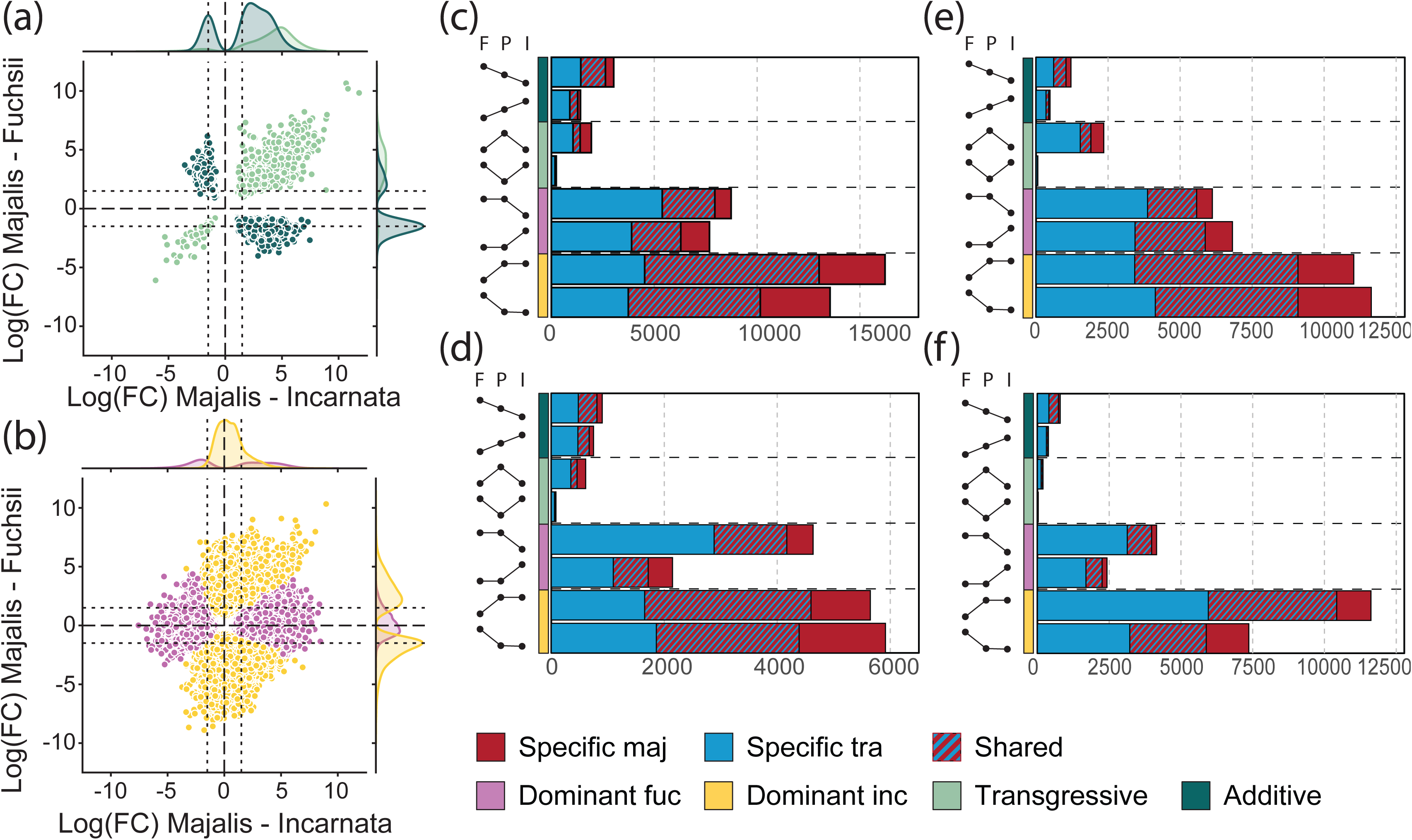
Patterns of genomic interactions in smRNA “targeting”. **(a–b)** Comparison of logFC values for *Dactylorhiza majalis* relative to either diploid parent (*D. fuchsii* on the Y-axis, *D. incarnata* on the X-axis) for all peaks (20–24 nt smRNAs) in annotated genes. **(a)** Colored symbols indicate additive and transgressive patterns; **(b)** shows dominance toward either diploid, as indicated in the legend. **(c-f)** Number of differentially targeted (DT) peaks in each type of genomic interaction, with peaks in *D. traunsteineri* shown in blue and *D. majalis* in red. Striped bars denote DT peaks shared between the allotetraploids. The annotation columns along the Y-axis represent the interaction type (colored bars, according to legend), while dots connected by lines show the direction of the interaction. Letters above each column indicate I - *D. incarnata*, P - polyploid, and F - *D. fuchsii.* **(c)** 20–24 nt smRNAs associated with genic regions, **(d)** 20–24 nt smRNA peaks outside annotated genic and TE regions, **(e)** 20–23 nt smRNAs in TEs, and **(f)** 24 nt smRNAs in TEs.

We also examined the distribution of interaction categories across genic regions (promoters, exons, introns). In *D. majalis*, nearly 75% promoter-associated DT peaks showed dominance towards *D. incarnata*, and almost 50% showed lower targeting than *D. fuchsii* (Supplementary Fig. S5). In contrast, in *D. traunsteineri* dominance toward *D. fuchsii* was more frequent across all genic regions (Supplementary Fig. S5).

Transgressively targeted genes in both allotetraploids were enriched for similar functional categories, including stress response, transport, transcription/translation regulation, and cell cycle control, though largely involving different genes and pathways in each species (Fig. 5b–c, Supplementary Tables S8–S9). In *D. traunsteineri*, transgressive peaks were associated with enrichment for cell cycle regulation and stress response functions, and the respective genes were targeted within exons or introns (Fig. 5b, Supplementary Tables S8– S9). In *D. majalis*, transgressive peaks occurred mostly within introns, and enrichments were associated with regulation of gene function (Fig. 5c, Supplementary Tables S8).

**Figure 5.**
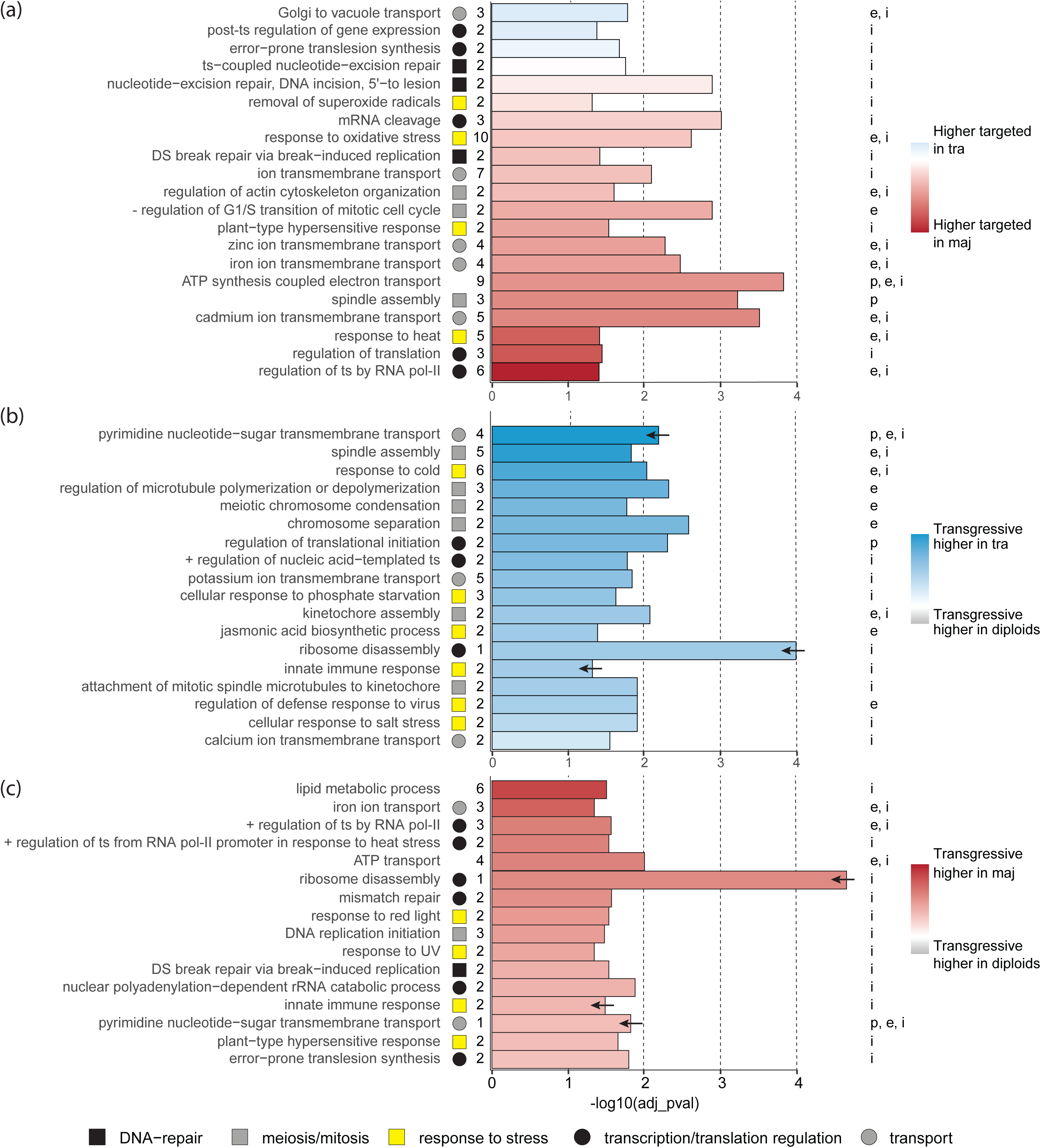
Summary of representative functional enrichments for **(a)** the allopolyploid comparison, **(b)** transgressive smRNA “targeting” of genic regions in *Dactylorhiza traunsteineri*, and **(c)** transgressive targeting in *D. majalis*. Bar length represents the adjusted significance of each enriched GO term. Full results are provided in Supplementary Tables S6, S8 and S9. Bar color indicates the z-score, computed from the logFC values of the corresponding targeting, as indicated in the legends. Arrows on bars in panels (b) and (c) mark GO terms shared between the two comparisons. Symbols next to the GO terms denote the broad functional categories (see legend), with numbers indicating the count of enriched genes within each category. Letters to the right of each plot specify the genetic region targeted by smRNAs; p - ‘promotor’, e - exon, i - intron. Abbreviations used in GO terms: + positive, - negative, DS double strand, pol polymerase, ts transcription.

### smRNAs associations with TEs

For TE-focused analyses, reads were divided into 24 nt smRNAs (likely het-siRNAs and therefore expected to be more frequently associated with TEs; Fig2c, Supplementary Fig. S1) and 20-23 nt smRNAs. After CPM filtering, for the comparative analysis of *D. majalis* versus *D. traunsteineri* more 24 nt peaks (315,838) passed the threshold than 20–23 nt peaks (102,222) (Supplementary Table S4). However, DT analysis uncovered the opposite trend: 3,242 DT peaks were identified for the 20–23 nt smRNAs, compared to only 103 DT peaks for the 24 nt smRNAs (Supplementary Table S4, Fig. 3d–e) suggesting a consistent targeting of TEs by het-siRNAs across allopolyploids. Very few TE-associated peaks were DT between the allotetraploids in both smRNA size classes simultaneously (Supplementary Fig. S6). In both datasets, approximately half of the DT peaks overlapped annotated TEs (Supplementary Table S4). Many of these were also located within gene introns, while a smaller number were found in the vicinity of gene promoters (Supplementary Table S4).

Similar to the results for the genic regions, *D. majalis* exhibited more frequent higher smRNA targeting of TEs than *D. traunsteineri* among DT peaks, except for 20–23 nt peaks associated with TEs located around promoters, where *D. traunsteineri* showed higher targeting (Supplementary Table S4). The majority of TE-associated DT peaks were located within LTR retrotransposons, especially Ty3-*gypsy* elements (Supplementary Fig. S7–S8). Among smRNAs targeting Ty3-*gypsy* elements, a slightly higher number targeted non-chromoviruses than chromoviruses (Supplementary Fig. S8).

Comparisons of allotetraploids versus diploids showed similar trends. After filtering, 139,736 (20-23 nt) and 370,440 (24 nt) peaks were retained (Supplementary Table S4). As with genic targets, DT peaks were classified into additive, transgressive, and dominance towards either diploid. In both datasets, a significant number of DT peaks were found to be shared between *D. majalis* and *D. traunsteineri* across interaction categories (p < 0.001, hypergeometric test; Fig 3d-e, Supplementary Fig. S3, Fig. 4e–f, Supplementary Fig. S7, Supplementary Fig. S9– 10).

Dominance patterns again predominated, with more peaks showing dominance towards either diploid than additive or transgressive patterns. For 20–23 nt smRNAs, slightly more peaks showed dominance toward the diploid with lower targeting, whereas for 24 nt smRNAs the opposite trend was observed (Fig. 4e–f, Supplementary Fig. S3, Supplementary Fig. S9). The contrast between dominant types (higher vs. lower targeting) was less pronounced for 20–23 nt but more distinct for 24 nt smRNAs (Fig. 4e–f, Supplementary Fig. S3, Supplementary Fig. S9–10). Three general patterns observed for smRNA-associated gene regions were also evident for TEs. First, for most additive peaks, the smaller genome of *D. fuchsii* exhibited higher smRNA targeting than the larger *D. incarnata* (Fig. 4e–f, Supplementary Fig. S3, Supplementary Fig. S9–10). Second, for transgressive peaks, both allotetraploids generally showed elevated smRNA targeting than either diploid (Fig. 4e–f, Supplementary Fig. S3, Supplementary Fig. S7, Supplementary Fig. S9–10). Finally, *D. traunsteineri* displayed greater differences from the parental diploids than *D. majalis* (Fig. 4e–f, Supplementary Fig. S3, Supplementary Fig. S7).

## DISCUSSION

smRNAs are key regulators of genome integrity and gene expression in polyploids, but most insights so far come from recent or resynthesized polyploids. In this study, we examine natural polyploids that have evolved over extended evolutionary timescales to provide a comprehensive overview of potential smRNA contributions to genomic and transcriptomic diversification following repeated allopolyploidization in *Dactylorhiza majalis* s.l. We analysed smRNA expression in two established sibling allotetraploids and their diploid progenitors, all grown under common garden conditions, thus minimizing environmental sources of variation. Our primary objectives were to comparatively characterize (*i*) patterns of genomic interaction in the allopolyploids and (*ii*) smRNA associations to TE control and gene activity.

Inferring smRNA *“targeting”* from sequence homology offers an informative yet indirect view of smRNA–genome interactions. Many plant smRNAs—particularly 24 nt siRNAs—act locally to reinforce transcriptional silencing of homologous TEs or nearby genes (Matzke & Mosher, 2014; Borges & Martienssen, 2015). Trans-acting and distal effects have nonetheless been documented (Lewsey *et al*., 2016; Liu & Chen, 2018), though the prevalence and biological impact of such long-distance effects remain less well characterized. Therefore, our analyses emphasize patterns of smRNA association rather than assuming direct functional regulation. These patterns highlight potential regions of epigenetic remodeling and provide a comparative framework for understanding smRNA landscape evolution after allopolyploidization.

We acknowledge that the present design—limited to representative individuals of the two extant diploid progenitors—cannot encompass the full spectrum of genetic and regulatory polymorphism that may have existed at the time of allopolyploid formation. Given that both parental lineages likely accumulated substantial variation over tens of thousands of generations, some of the observed differences between the allopolyploids may partly reflect ancestral or subsequent divergence in the diploid gene pools. Accordingly, we interpret our results in terms of general and reproducible regulatory patterns rather than exact reconstructions of the parental regulatory states.

### Consistent dominance in the allopolyploids towards the paternal, larger genome

Allopolyploidization merges divergent genomes, giving rise to novel genetic and regulatory interactions. Common outcomes include epistasis and non-additive genetic or epigenetic effects, including smRNA-mediated gene regulation and TE silencing. Our results reveal a consistent dominance of smRNA targeting aligned to the paternal, larger subgenome (*D. incarnata*), especially for TE-associated 24 nt smRNAs. This pattern was observed across both sibling allotetraploids (Fig. 4, Supplementary Fig. S5), and resulted in a comparatively small number of DT peaks between them. Similar non-additive smRNA responses following hybridization and/or whole-genome duplication have been documented in *Arabidopsis* (Ha *et al*., 2009), *Brassica* (Fu *et al*., 2016; Martinez Palacios *et al*., 2019), and *Spartina* (Cavé-Radet *et al*., 2020), though it has remained unclear whether such patterns remain consistent across longer evolutionary timescales.

Examining whether smRNA expression is additive, dominant, or transgressive allows us to understand how regulatory interactions from the parental genomes combine in allopolyploids. Additive patterns indicate intermediate regulation and minimal epistatic interactions, dominance reflects the stronger influence of one parental subgenome, and transgressive patterns reveal novel regulatory states beyond the parental range. These distinctions are conceptually important because they highlight mechanisms of genome stabilization, resolution of redundant gene copies, and the generation of potentially adaptive phenotypes that can enable allopolyploids to occupy ecological niches outside the parental range (Ha et al., 2009; Fu et al., 2016; Chen et al., 2020).

Given that *D. incarnata* has a larger genome and higher TE load than *D. fuchsii* (Eriksson *et al*., 2022), our findings are consistent with prior reports showing that smRNA-mediated silencing preferentially targets the TE-rich subgenome, as observed in wheat (Woodhouse *et al*., 2014) and *Brassica rapa* (Cheng *et al*., 2016). In contrast, gene expression dominance in allopolyploids frequently shifts toward the subgenome with fewer TEs, a pattern that can arise rapidly after allopolyploidization (for example in *Mimulus*; Edger *et al*., 2017) and persist across generations (Gaebelein *et al*., 2019; Liu & Wang, 2023).

While our data indicate stronger dominance towards *D. incarnata*, some patterns may be influenced by reference genome bias, as we only had access to a *D. incarnata* reference genome. Accurate differentiation between homologous and homeologous targeting remains challenging due to the short length of smRNAs and their tolerance for mismatches in plants. Nonetheless, mapping success was comparable across samples (ca. 83%; Supplementary Table S3), and intergenic smRNA dominance towards *D. fuchsii* was also observed, suggesting that reference bias alone cannot account for our observations. Lineage-specific TE degradation or divergence in *D. fuchsii* may have reduced smRNA detection from this genome. Notably, *D. incarnata* and *D. fuchsii* diverged approximately 5.5 million years ago (Brandrud *et al*., 2020).

### Divergent targeting of TEs by 24 nt and 20-23 nt smRNAs

As expected, 24 nt smRNAs—typically het-siRNAs—were strongly associated with TEs, particularly LTR retrotransposons (Fig. 2c, Supplementary Fig. S1). Surprisingly, the number of DT peaks for 24 nt smRNAs between the allotetraploids was low, suggesting that TE silencing proceeds in a relatively consistent manner across these independently evolved lineages (Fig.3e). In contrast, 20–23 nt smRNAs exhibited markedly greater divergence between the sibling allopolyploids (Fig. 3d), consistent with a more dynamic, lineage-specific regulatory role.

Previous studies have shown that TE-associated smRNA activity often peaks shortly after hybridization or whole-genome duplication, followed by stabilization via DNA methylation or other epigenetic mechanisms (Jiao *et al*., 2018; Martinez Palacios *et al*., 2019). In *Arabidopsis*, Ha *et al*., (2009) proposed that allopolyploids exploit smRNAs inherited from the parents to buffer genomic shock and promote rapid stabilization. The comparatively low divergence in 24 nt smRNA targeting observed here may reflect the advanced age of these natural allotetraploids—approximately 74k (*D. traunsteineri*) and 104k generations (*D. majalis*) (Hawranek, 2021). Nonetheless, for the few DT peaks detected, smRNA abundance was higher in the older *D. majalis* (Fig. 3), possibly indicating continued regulatory refinement associated with its broader ecological range or a more advanced stage of genome diploidization.

Intriguingly, most smRNA differences associated with TE regulation in the ecologically divergent allotetraploids involved 20–23 nt smRNAs, which likely include miRNAs and nat-siRNAs. These classes are frequently implicated in responses to biotic and abiotic stress (Ito *et al*., 2011; Marí-Ordóñez *et al*., 2013; Creasey *et al*., 2014; Lee *et al*., 2020). Elevated expression of miRNAs (typically 20–22 nt long) has also been observed in resynthesized allopolyploids relative to diploid progenitors across several systems (Ha *et al*., 2009; Kenan-Eichler *et al*., 2011; Ghani *et al*., 2014), underscoring the potential role of shorter smRNAs in stress adaptation and dynamic gene regulation in polyploids.

Despite *Ty1-copia* being the most abundant LTR-TE in *Dactylorhiza* genomes (Eriksson *et al*., 2022), we observed more 20–23 nt smRNAs mapping to, and possibly derived from, *Ty3-gypsy* elements (Supplementary Figs. S6–S7), particularly non-chromovirus subgroups. Notably, *Ty3-gypsy* elements displayed contrasting patterns between the diploid progenitors: chromovirus subgroups were more prevalent in *D. fuchsii*, while non-chromoviruses were more common in *D. incarnata* (Eriksson *et al*., 2022). Elevated targeting of *Ty3-gypsy* elements by shorter smRNAs in allopolyploids may reflect subgenome conflict or TE activation following hybridization. A similar phenomenon occurs in *Gossypium*, where TEs originally in the higher-TE A-subgenome transposed into the D-subgenome post-allopolyploidization (Chen *et al*., 2020), suggesting that transposition between subgenomes may contribute to regulatory instability and diversification.

### Genic smRNA targeting and possible regulatory implications

Polyploid formation introduces both meiotic/mitotic challenges and extensive genetic redundancy. Accurate differentiation between homologous and homeologous chromosomes is essential to ensure correct meiotic segregation. Preferential pairing of homologs has been observed in early stages of polyploid formation in *Arabidopsis suecica* (Pecinka *et al*., 2011) and *Brassica napus* (Grandont *et al*., 2014), although elevated homeolog mispairings has also been reported (Xiong *et al*., 2021). Simultaneously, allopolyploids often undergo genetic diploidization, in which some redundant or misregulated gene copies may be silenced or lose function over time (Lee & Chen, 2001; Kashkush *et al*., 2002; Koh *et al*., 2010). Some duplicated copies can be retained and remain functional, for example if they confer adaptive benefits, become subfunctionalized, or acquire new functions via neofunctionalization (Adams *et al*., 2003; Blanc & Wolfe, 2004; Duarte *et al*., 2006; De Smet *et al*., 2013).

In addition to silencing TEs, smRNAs—particularly 21–22 nt species—regulate gene expression via PTGS or TGS (Borges & Martienssen, 2015). We observed widespread genic smRNA targeting, especially in exons and 1 kb upstream of gene models (putative promoters), with 21 nt smRNAs contributing the most. While such targeting may mediate transcript degradation, alternative splicing, or cis-silencing (Guleria *et al*., 2011; Moss, 2001), some associations may also have no functional consequence. Nevertheless, these patterns can reflect regulatory adjustments during early diploidization.

Substantial genic smRNA targeting was also detected in the diploids, especially *D. fuchsii*, likely reflecting stress responses in the common garden experiment conducted outdoors in Vienna, Austria—an area where only *D. incarnata* occurs naturally. Environmental mismatch may have triggered miRNA- and nat-siRNA-mediated regulation in *D. fuchsii*, consistent with previous reports of stress-induced smRNA upregulation (Zhang *et al*., 2012; Mao *et al*., 2021; Tiwari & Rajam, 2022).

Transgressive smRNA expression relative to the diploid progenitors may reflect how allopolyploids respond to genome duplication while also acquiring novel ecological features, potentially facilitating establishment in niches outside the parental range. We observed more transgressive activity at peaks targeted by 20–23 nt smRNAs—compared to 24 nt— especially when the allotetraploids showed higher smRNA levels relative to both diploid parental species (Fig. 4e–f).

Genes transgressively targeted by smRNAs were enriched for functions related to mitosis, meiosis, gene regulation, stress responses, and transport (Fig. 5). While these functional categories were largely shared across the two allotetraploids, the specific genes and pathways involved differed, suggesting species-specific regulatory routes. In the younger *D. traunsteineri*, enrichment of smRNA targeting was observed for genes involved in cell cycle control, possibly reflecting ongoing meiotic stabilization. By contrast, the older *D. majalis* showed enrichments for genes involved in transcriptional and translational regulation, suggesting a later phase of polyploid evolution focused on dosage control and fine-tuning of gene expression.

Together, these observations support a dynamic evolution of smRNA-mediated regulation following allopolyploidization: an early phase dominated by stabilization of chromosome behavior during meiosis, together with refinement of gene redundancy and the emergence of functional divergence. Similar dynamics have been proposed in other systems including *Arabidopsis*, *Brassica*, and *Gossypium* (Ha *et al*., 2009; Fu *et al*., 2016; Chen *et al*., 2020).

### Conclusions and Future Directions

This study underscores the dual associations of smRNAs with genomic features in allopolyploids—both with TEs and genic regions—reflecting their potential roles in genome regulation. Despite their shared ancestry, *D. majalis* and *D. traunsteineri* exhibit clear lineage-specific patterns of smRNA activity, particularly in genic regions, suggesting divergent regulatory trajectories following allopolyploidization. Given their rapid action, diversity, and regulatory plasticity, smRNAs likely play central roles in both early and long-term genomic reorganization of allopolyploids. Our results support a model in which smRNAs mediate both immediate stabilization and longer-term functional divergence after genome merging. Future work should integrate smRNA expression profiles with homeolog-specific reference genomes and DNA methylation landscapes. These approaches would provide deeper insights into subgenome-specific regulation, the dynamics of smRNA–target interactions, and the evolutionary consequences of smRNA-mediated gene and TE regulation in natural allopolyploids.

## ACKNOWLEDGEMENTS

This work was funded by the Austrian Science Fund (FWF) [grant DOI: 10.55776/Y661 awarded to OP and through the doctoral programme DOI: 10.55776/W1225, awarded to a faculty team including OP]. For open access purposes, the author has applied a CC BY public copyright license to any author accepted manuscript version arising from this submission. Faculty members, in particular Andreas Futschik, Robert Kofler and Christian Schlötterer, as well as the students and the scientific advisory board of the Vienna Graduate School of Population Genetics (www.popgen-vienna.at), Richard Bateman, and Ortrun Mittelsten Scheid are gratefully acknowledged for their valuable discussions and feedback throughout this work. We thank Juliane Baar, Daniela Paun and Maria Teresa Lorenzo for their support with laboratory procedures. Sequencing was performed at the Vienna BioCenter CoreFacilities (VBCF; https://www.viennabiocenter.org/). Computational resources were provided by the Life Science Compute Cluster (LiSC) of the University of Vienna. We also acknowledge Natural England and the Forestry Commission (UK), and regional county administrations in Austria, France, and Sweden for issuing the necessary collecting permits.

## AUTHOR CONTRIBUTIONS

Study conceived and designed by OP. Sampling and transplantation to the common garden conducted by OP, FB, MH, MCE and TMW. Read processing and mapping done by MT. Identification of putative smRNA peaks across the genome performed by MT and MCE. Data analyses of differentially targeted genes and TEs conducted by MCE. Analysis of overlapping differentially expressed and differentially targeted genes done by TMW. Interpretation of the results was undertaken by MCE. The manuscript was drafted by MCE with feedback from OP, and was revised and approved by all authors.

## DATA AVAILABILITY

Newly reported raw Illumina smRNA-seq data are deposited on NCBI SRA (GenBank BioProject PRJNA317244, accessions SRR17191877–SRR17191897). Previously published Illumina RNA-seq and smRNA-seq are available from the same GenBank BioProject PRJNA317244. The pipeline, scripts and raw count file, together with the peak gff3 file are available at https://github.com/mc-er/dact-sRNAs.

