## Supplementary Figures for "Small RNAs regulation and genomic harmony: insights into allopolyploid evolution in marsh orchids (*Dactylorhiza*)"

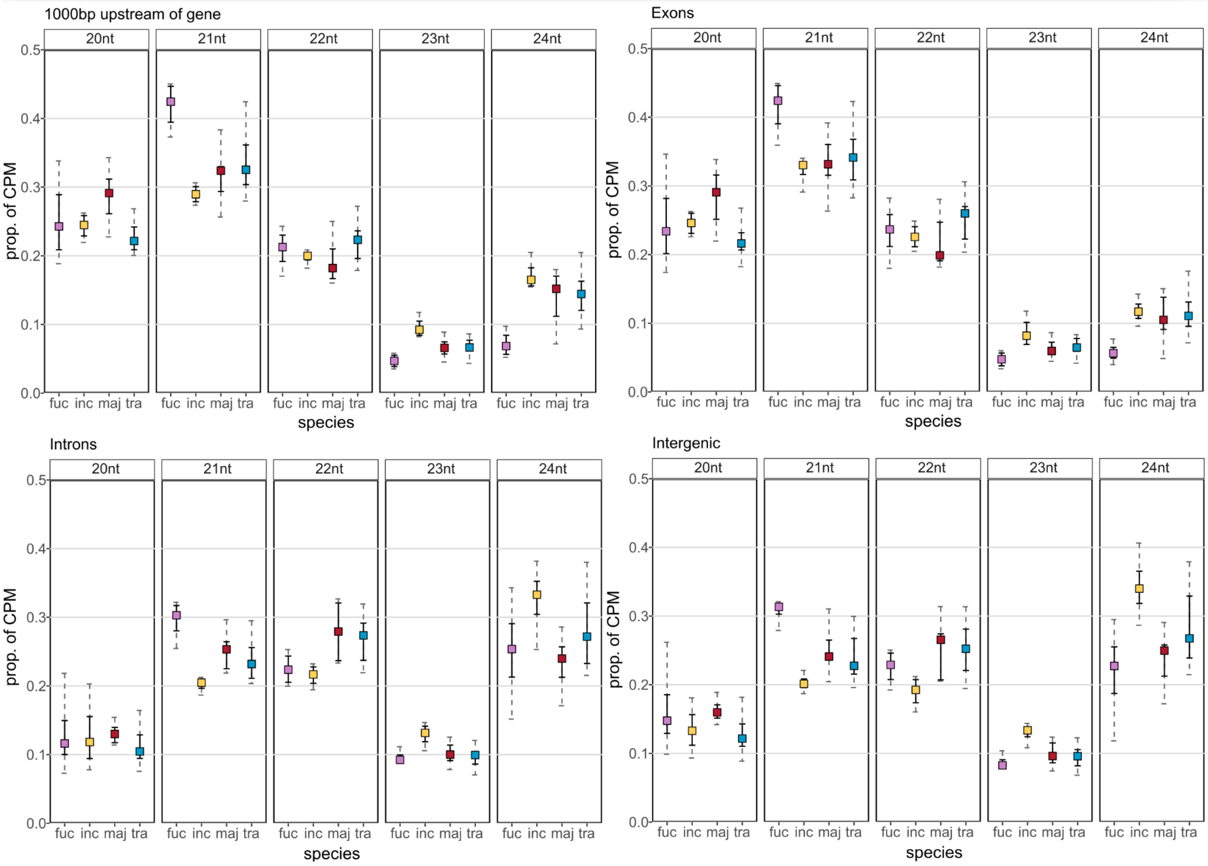


**Figure S1.** Distribution of normalised reads counts for different smRNA lengths over genomic regions (promoter/1000 bp upstream, exons, introns and intergenic) in the four tested species. fuc = *D. fuchsii*, inc = *D. incarnata*, maj = *D. majalis*, tra = *D. traunsteineri*.

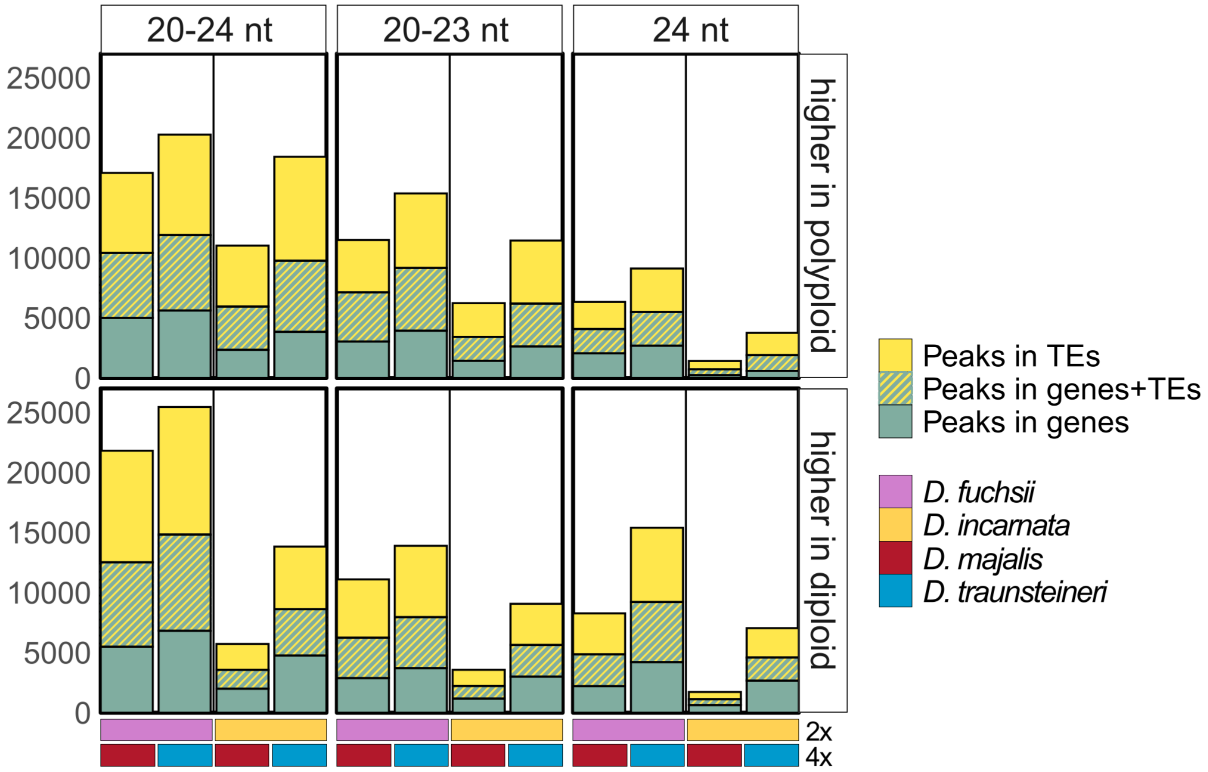


**Figure S2.** Number of differentially targeted peaks in genes and TEs for each comparison between a polyploid (4x) and a diploid (2x) for all three data sets, 20-24 nt 20-23 nt and 24 nt smRNAs. The top panel contains the number of peaks where the polyploid is over regulated compared to the diploid, whereas the bottom panel contains the number of peaks where the diploid is over targeted. The coloured boxed below the figure represents the comparison shown in the corresponding column.

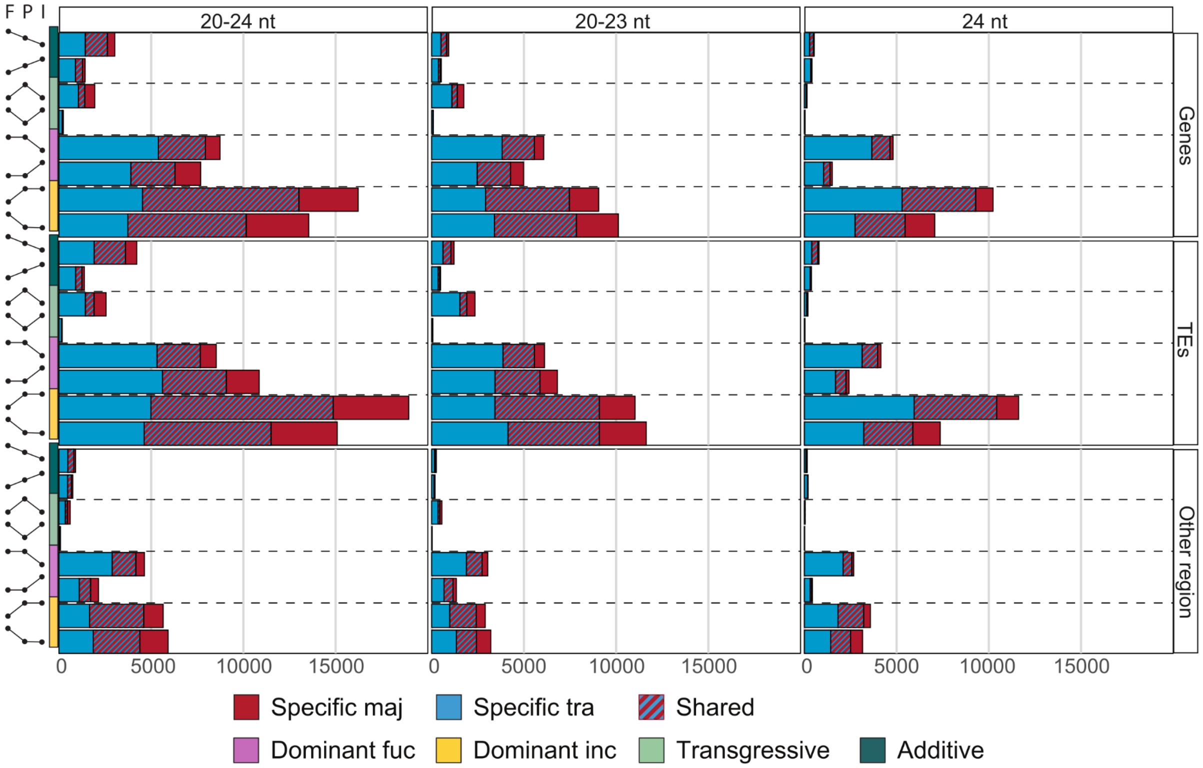


**Figure S3.** Number of DT peaks in all datasets 20−24 nt, 20−23 nt and 24 nt smRNAs for each genomic interaction (additive, transgressing or dominant to either diploid). The top row contains DT peaks in genes, the middle row shows DT peaks in TEs and the bottom row shows peaks found outside annotated genic and TE regions, e.g. intergenic. Colours represent, red - *D. majalis*, blue - *D. traunsteineri* and striped pattern - number of DT peaks showing the same genomic interaction in both *D. majalis* and *D. traunsteineri*. The two annotation columns along the Y−axis represent the type (coloured bars according to legend) of genomic interaction Fig. 4, while dots connected with lines show the direction. Letters above each column of dots represent **F** - *D. fuchsii*, **P** - polyploid and **I** - *D. incarnata*.

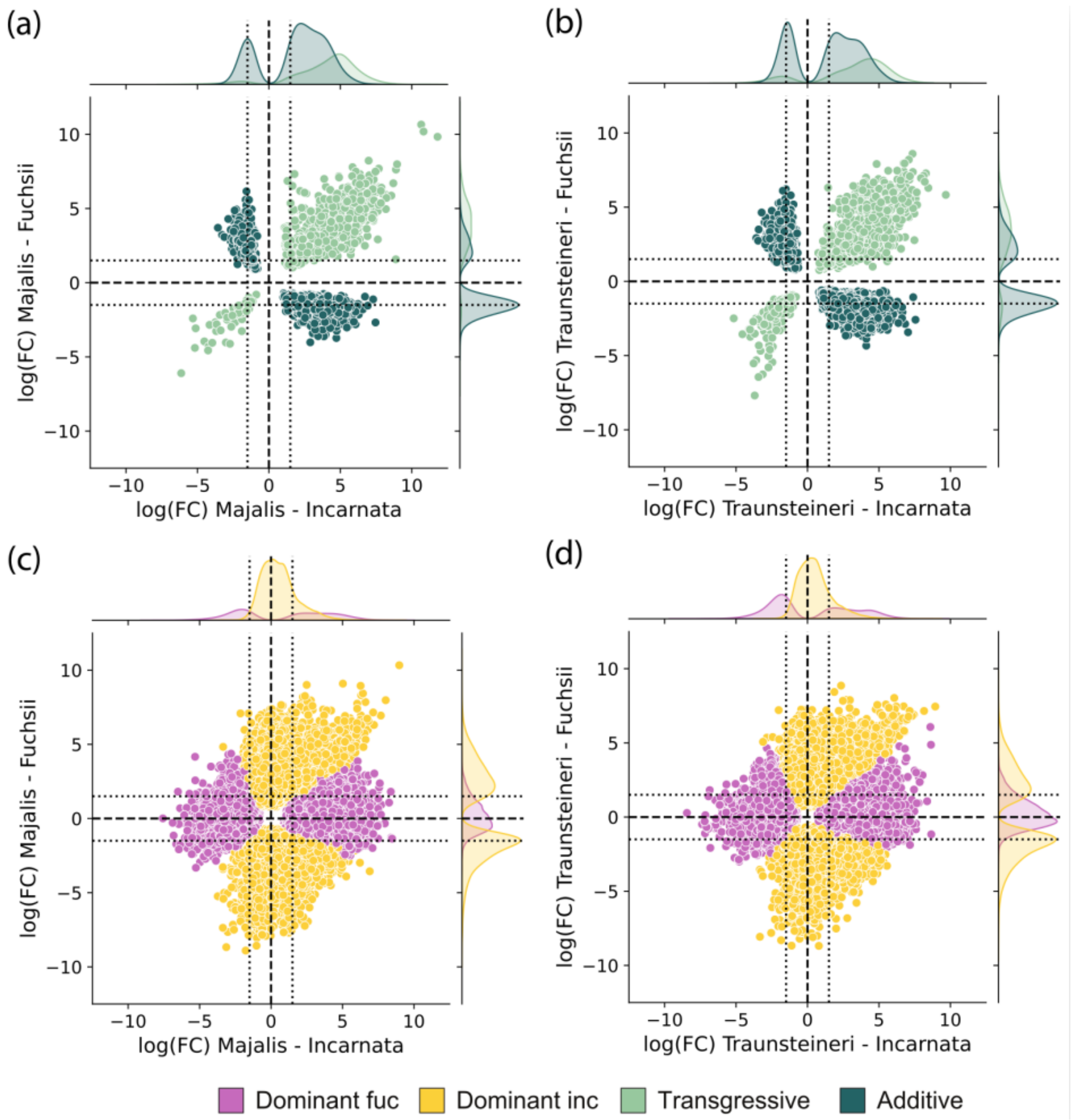


**Figure S4.** Patterns of genomic interactions comparing logFC values for *D. majalis* (left) and *D. traunsteineri* (right) towards either diploid, *D. fuchsii* on the Y-axis and *D. incarnata* on the X-axis for all peaks (i.e., 20−24 nt smRNAs) found in annotated genes. (a-b) shows with coloured symbols transgressive and additive patterns; whereas (c-d) is dominant to either diploid, according to the legend.

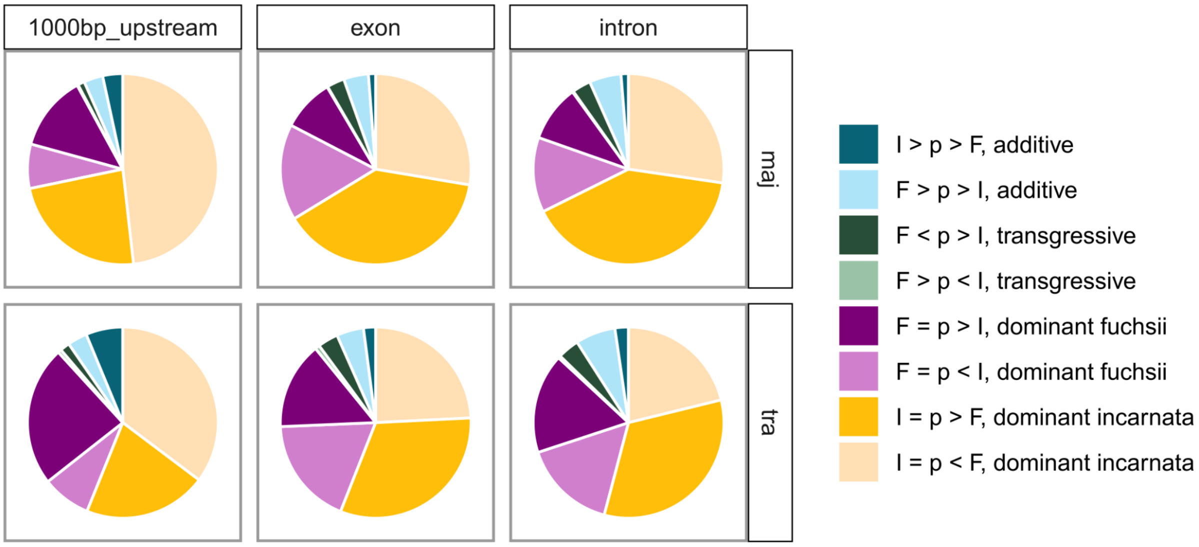


**Figure S5** Proportions of peaks within each genomic interaction (additive, transgressing or dominant to either diploid) for each genomic region (1000 bp upstream/promoter, exon, intron). Colours are according to legend. Letters in legend represent **F** - *D. fuchsii*, **p** - polyploid and **I** - *D. incarnata* while symbols represent level of smRNA between the diploids and polyploid.

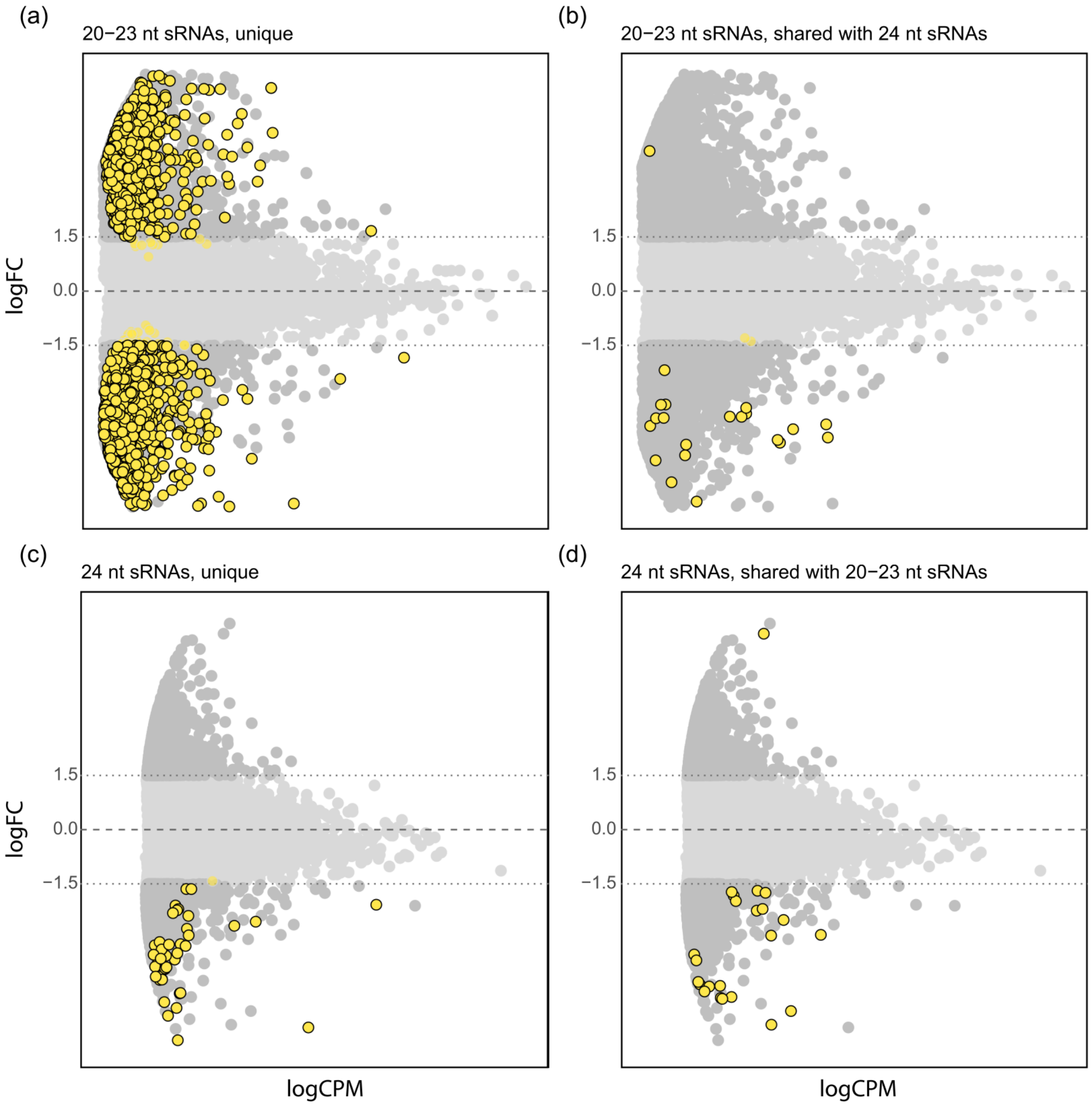


**Figure S6** MA plots showing differences between *D. majalis* and *D. traunsteineri* for TEs types grouped together, the central dashed line marks 0 while the dotted lines mark logFC -1.5 and 1.5. Negative logFC values represent over-targeting in *D. majalis* as compared to *D. traunsteineri*, whereas positive logFC values indicate over-targeting in *D. traunsteineri*. (a) and (b) show 20−23 nt smRNAs while (c) and (d) show 24 nt smRNAs. (a) and (c) show DT peaks specific for each smRNA group, i.e. 20−23 nt or 24 nt, while (b) and (d) show DT peaks for both 20−23 nt and 24 nt smRNAs.

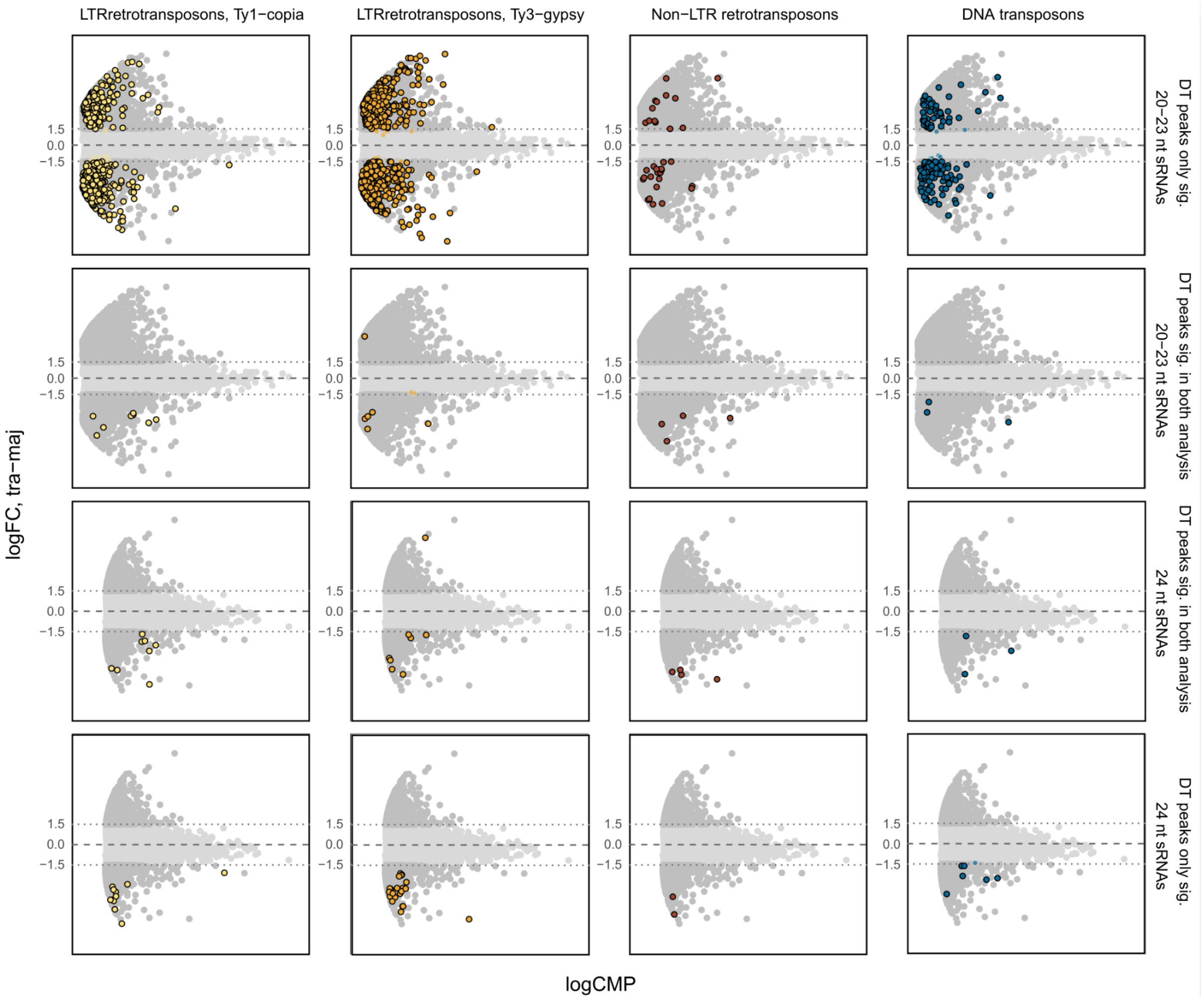


**Figure S7** MA plots showing differences between *D. majalis* and *D. traunsteineri* for TEs, the central dashed line marks 0 while the dotted lines mark logFC -1.5 and 1.5. Negative logFC values represent over-targeting in *D. majalis* as compared to *D. traunsteineri*, whereas positive logFC values indicate over-targeting in *D. traunsteineri*. Colours correspond to different types of TEs, indicated by column headers. Rows correspond to smRNAs, where the 1st and 2nd row show 20−23 nt smRNAs and 3rd and 4th show 24 nt smRNAs. The 2nd and 3rd row show peaks that are DT for both 20−23 nt and 24 nt smRNAs.

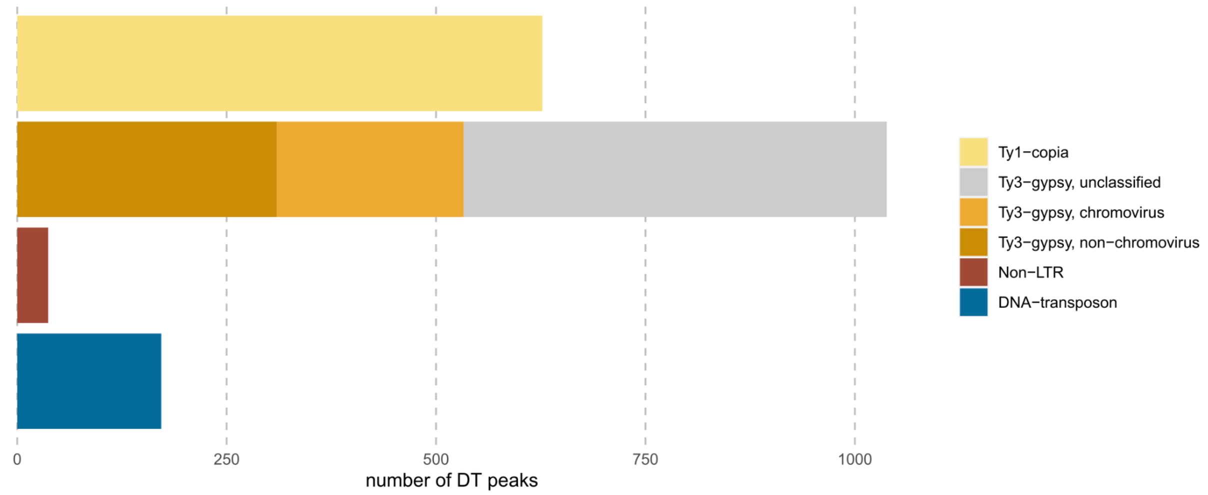


**Figure S8** Count of DT peaks between *D. majalis* and *D. traunsteineri*, the coloured points of the plots in the 1st row (i.e. 20−23 nt smRNAs) in Fig. S7. Here Ty3-gypsy has been divided into the next annotation level.

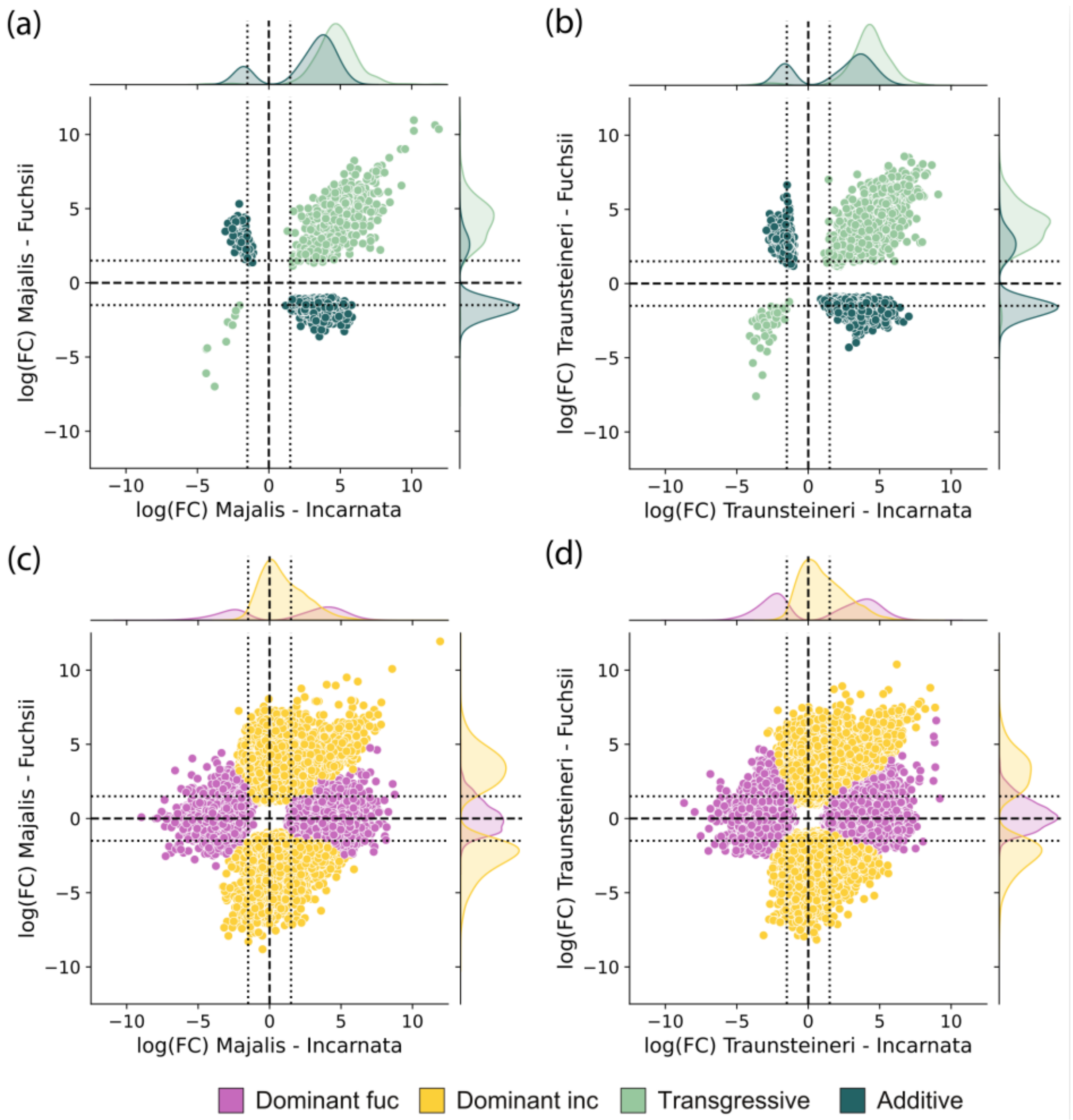


**Figure S9** Patterns of genomic interactions comparing logFC values for *D. majalis* (left) and *D. traunsteineri* (right) towards either diploid, *D. fuchsii* on the Y-axis and *D. incarnata* on the X-axis for 20-23 nt smRNAs peaks found in annotated TEs. Top row shows with coloured symbols transgressive and additive patterns; whereas bottom row shows dominant to either diploid.

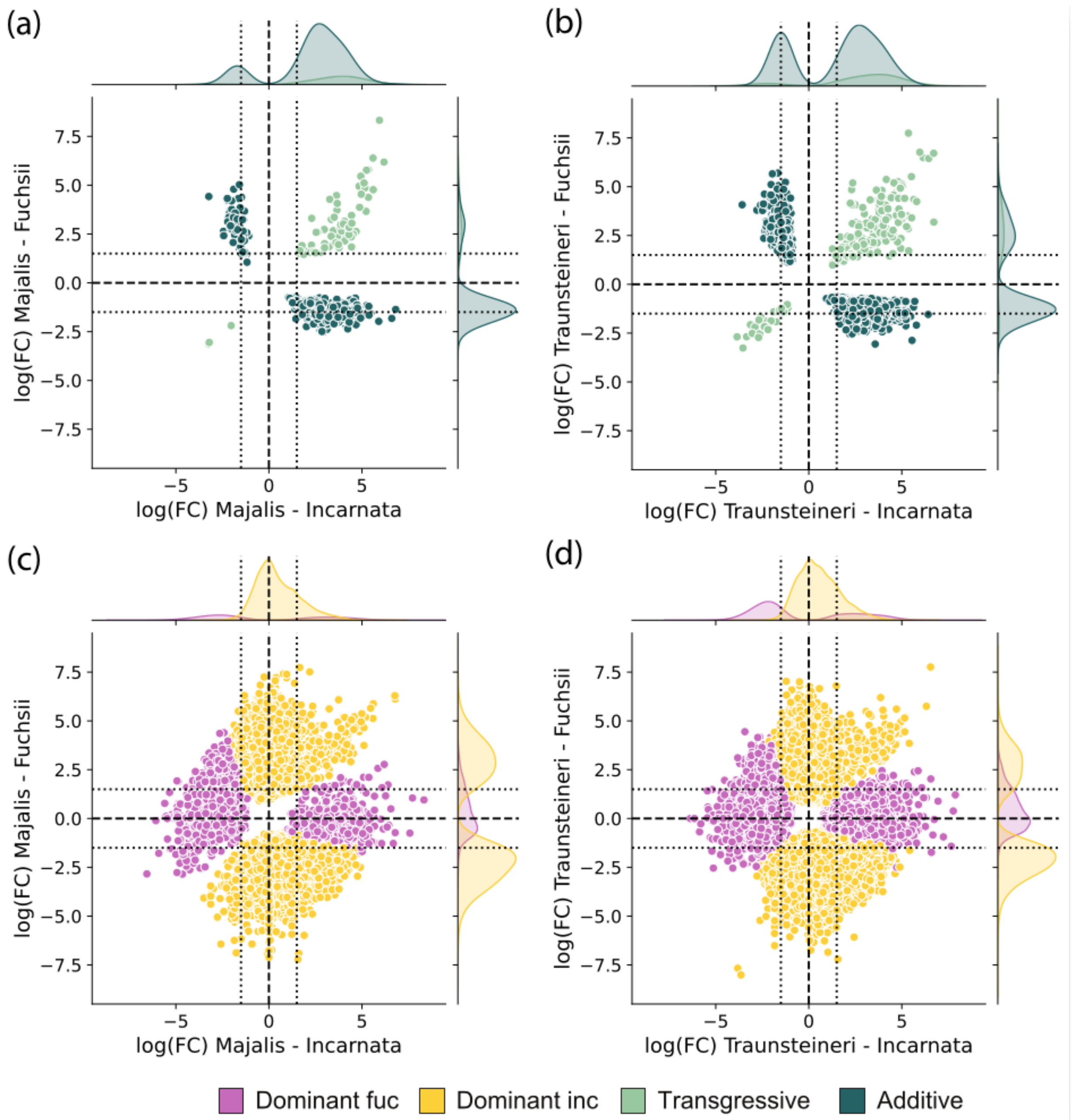


**Figure S10** Patterns of genomic interactions comparing logFC values for *D. majalis* (left) and *D. traunsteineri* (right) towards either diploid, *D. fuchsii* on the Y-axis and *D. incarnata* on the X-axis for 24 nt smRNAs peaks found in annotated TEs. Top row shows with coloured symbols transgressive and additive patterns; whereas bottom row shows dominant to either diploid.
